# Raman imaging and statistical methods for analysis various type of human brain tumor

**DOI:** 10.1101/2021.04.08.438923

**Authors:** M. Kopec, M. Błaszczyk, M. Radek, H. Abramczyk

## Abstract

Spectroscopic methods provide information on the spatial localization of biochemical components based on the analysis of vibrational spectra. Raman spectroscopy and Raman imaging can be used to analyze various types of human brain tissue. The objective of this study is employment of Raman spectroscopy and Raman imaging to evaluate the Raman biomarkers to distinguish tumor types. We have demonstrated that bands characteristic for carotenoids (1156 cm^−1^, 1520 cm^−1^), proteins (1004 cm^−1^), fatty acids (1444 cm^−1^, 1655 cm^−1^) and cytochrome (1585 cm^−1^) can be used as universal biomarkers to distinguish aggressiveness in human brain tumor. The sensitivity and specificity obtained from PLS-DA have been over 85%. Only for pituitary adenoma the specificity is lower and takes equal 46%. The presented results confirm the potential applications of vibrational spectroscopy methods in oncological diagnostics.

## Introduction

The nervous system tumors are one of the most common types of cancer in the world. ^1^ In this paper we analyzed various types of human brain tumor such as: gliosarcoma ^2^, meningothelioma, anaplastic oligodendroglioma ^3^, pituitary adenoma ^4^, neurofibroma, testicular cancer metastasis to the brain and breast cancer. ^5^

According to WHO meningotheliomas (Meningioma menigotheliale) account for some 15-18% of all intracranial tumors. Meningotheliomas are the most frequent brain tumors in adults and they occur more often in female than in male. ^6^ When meningotheliomas are in benign stadium they carry a good prognosis. Standard treatment for meningotheliomas is surgical resection. It is estimated that 80% of meningotheliomas are grade I (mildly malignant). ^7^ The main symptoms in patient with meningotheliomas are sensory and motor deficits or gait disturbance.^8^

Malignant glioma is the most common type of primary brain tumor. There are known three subtypes of gliomas such as: anaplastic astrocytoma (AA), anaplastic oligoastrocytoma (AOA), anaplastic oligodendroglioma (AO), and anaplastic ependymoma. Anaplastic oligodendroglioma tumors are rare and uncommon; they constitute 2-7% of primary brain tumors. ^9^ The standard therapy for patients with anaplastic oligodendroglioma and the treatment include a lot of therapies such as: chemotherapy ^10^, stereotactic radiotherapies ^11^, targeted therapy ^12^, and reoperation. ^13^ Despite intensive study, the mortality among patients with malignant glioma is very high. It is estimate that 77 % die within 1 year after diagnosis.^14^

A neurofibroma is a type of nerve tumor. Neurofibromas usually are benign peripheral nerve sheath usually single and commonly occur in all parts of the body. A neurofibroma may appear in people with a genetic disorder (neurofibromatosis type 1) or can arise with unknown cause. These nerve tumors most often occur in people of age 20 to 40 years. Neurofibromas look like little “rubber balls” under the skin. They can also protrude from the skin. ^15^ For the first time the neurofibroma was described by Von Recklinghausen in 1882. ^16^ The morbidity and mortality caused by neurofibromatosis depend on the occurrence of complications that involve any of the body system.^17^

Pituitary adenomas are tumors occurring in the anterior pituitary. Most pituitary tumors are benign and slow-growing. Pituitary adenoma can be classified as microadenoma, macroadenoma, and giant tumors depending on the tumor size. Tumors smaller than 10 mm are considered to be microadenomas, while macroadenomas are tumors larger than 10 mm. In contrast giant pituitary tumors are larger than 40 mm. ^18^ For patients with pituitary adenomas there are available treatment options such as: surgical excision, medical and radiation therapy.^18^

When cancer cells spread from their original site to the brain, metastases occur. The most likely types of cancer which spread to brain are: lung cancer, melanoma, colon cancer and breast cancer, less often testicular cancer. It is estimated that brain metastases occur in 10 to 30 percent of adults with diagnosed cancer. ^19^

Surgery is the most common form of treatment for brain tumor.

One of the main reasons for insufficient progress in brain tumor diagnostics is related to the fact that most cancer types are not only heterogeneous in their genetic and biochemical composition but also reside in varying microenvironments and interact with different cell types. Until now, no technology has been fully proven for effective detecting of invasive cancer, infiltrating the extracellular matrix. So that, development of novel treatment strategies is needed. Tashibu presented first results obtained with the use of Raman spectroscopy in brain tumor analysis. ^20^ He analyzed the water content in brain tissues. This preliminary study gave impetus to further investigations. Using Raman spectroscopy and Raman imaging made possible showing the distribution of biochemical components in brain tumors. Those components can be divided into subgroups such as: structural ^21^, metabolic ^22^, epigenetic ^23^, immunologic ^24^ and genetic ^25^.

In this paper we present Raman spectroscopy as a tool for distinguishing various subtypes of human brain tumor.

## Results and discussion

The experiments were carried out using Raman spectroscopy and Raman imaging. Firstly, we focused on the Raman spectroscopy analysis for mildly malignant and malignant brain tumors.

Figure 1 present the video image, Raman image and Raman spectra of the tissue of meningothelioma (Meningioma meningotheliale, WHO grade I). Raman images were performed by using Cluster Analysis method in the spectral region 400-3300 cm^−1^.

**Figure 1.**
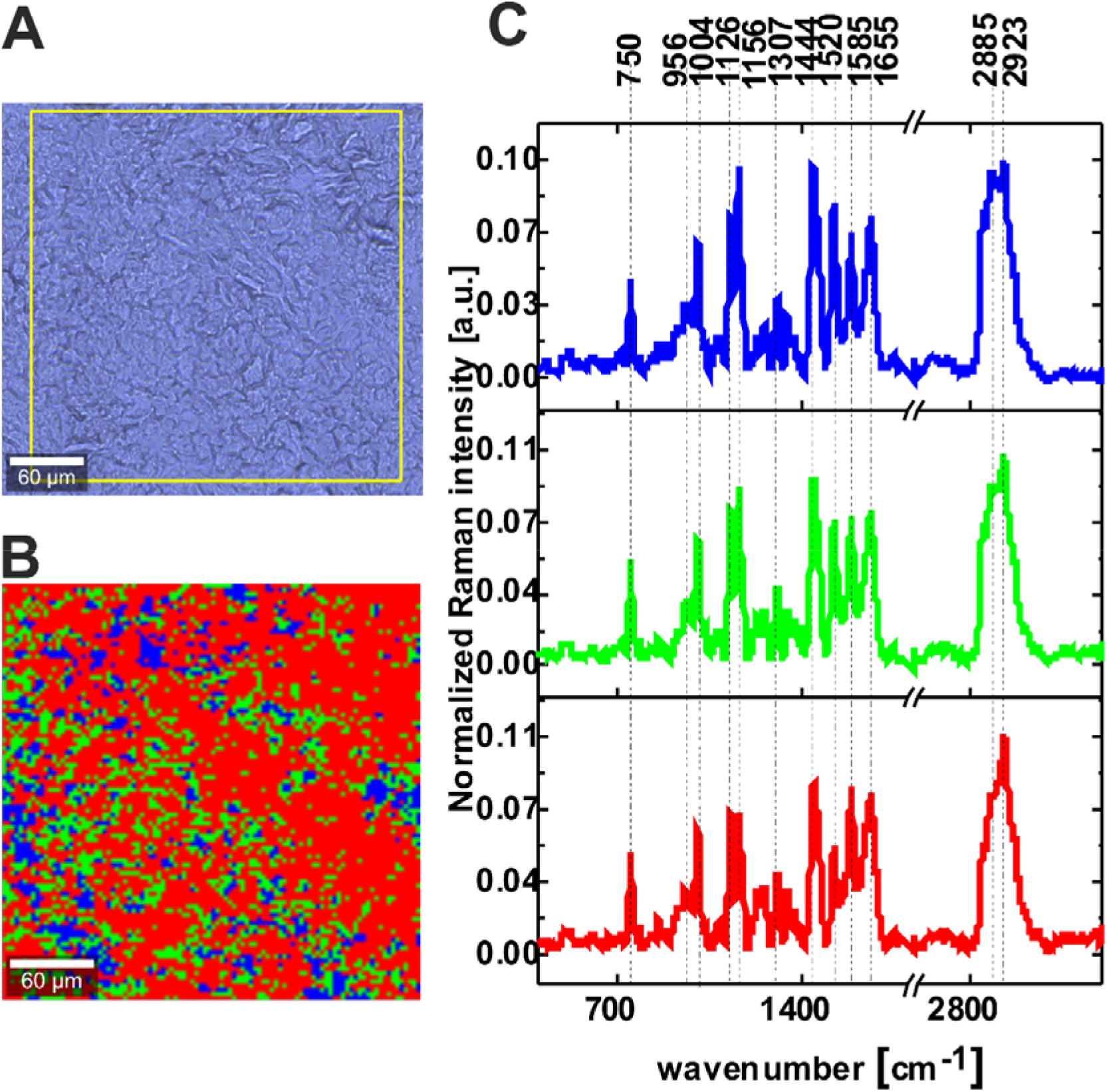
*Microscopy image (A), Raman image (300 µm x 300 µm) (B) and Raman spectra (C) obtained by Cluster Analysis method in range 400-3300 cm^−1^ from meningothelioma WHO grade I). Integration time for Raman images 0.5 s in the high frequency region and 1s in the fingerprint region. The colours of the lines of the Raman spectra correspond to the colours of Raman image*.

From Figure 1 we can see that spectra of mildly tumors are dominated by the peaks at 750, 956, 1004, 1126, 1156, 1307, 1444, 1520, 1585, 1655 cm^−1^ in fingerprint region and by the peaks at 2885, 2923 cm^−1^ in high frequency region. Detailed spectral information for biochemical components assigned to the Raman vibrations are presented in Table 1.

**Table 1.**

| Frequency [$\text{cm}^{-1}$] | Assignment |
| --- | --- |
| 750 | cytochrome |
| 956 | hydroxyproline/Collagen backbone |
| 1004 | proteins, phenylalanine |
| 1126 | cytochrome |
| 1156 | carotenoids |
| 1228 | nucleic acids (Try, Ala)/Proteins (Amide III $\beta$ sheet or random |
|  | coil),<br>Lipid, phospholipid =C-H bend |
| 1307 | lipids, phospholipids<br>CH<sub>2</sub> twist, collagen, protein amide III, DNA |
| 1444 | fatty acids, triglycerides, CH<sub>2</sub> or CH<sub>3</sub> |
| 1520 | carotenoids |
| 1585 | cytochrome |
| 1655 | unsaturated fatty acids, triglycerides (C=C) str, amide I $\alpha$ helix |
| 2885 | lipids, =CH str |
| 2923 | proteins |
| 3164 | str (NH), str (OH), amide, water |

To compare the biochemical composition of mildly malignant tumor brain tissue we conducted the same analysis for malignant brain tissue (anaplastic oligodendroglioma WHO grade III). Figure 2 presents the video image, Raman image and Raman spectra of the tumors brain tissue.

**Figure 2.**
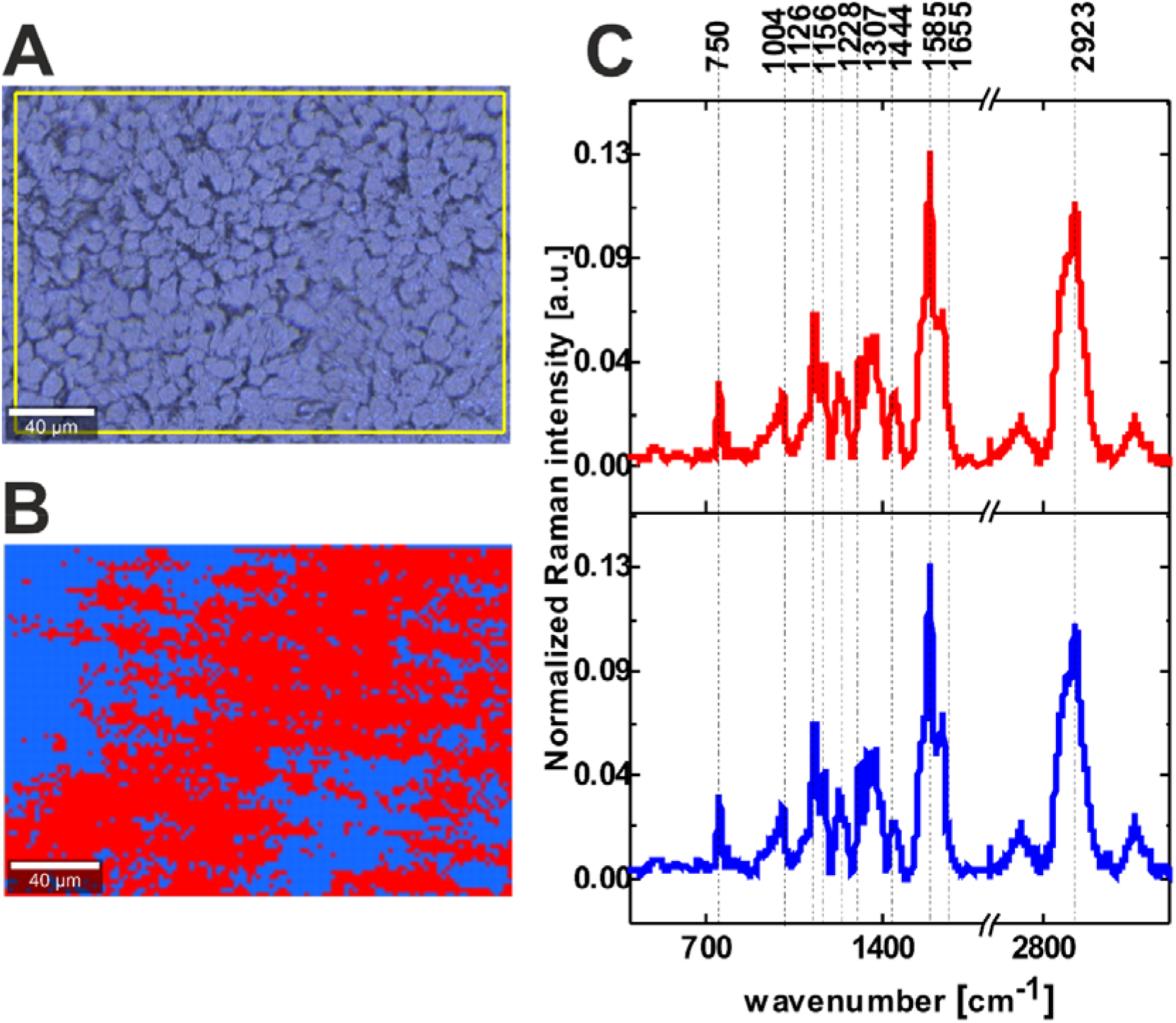
*Microscopy image (A), Raman image (230 µm x 160 µm) (B) and Raman spectra (C) obtained by Cluster Analysis methods in range 400-3300 cm^−1^ anaplastic oligodendroglioma WHO grade III). Integration time for Raman images 0.5 s in the high frequency region and 1s in the fingerprint region. The line colours of the Raman spectra correspond to the colours of Raman image.*

From Figure 2 we can see that spectra of malignant human brain tumor are dominated by the peaks at 750, 1004, 1126, 1156, 1228, 1307, 1444, 1585, 1655 cm^−1^ in fingerprint region and by the peak at 2923 cm^−1^ in high frequency region.

Detailed spectral information for biochemical components are presented in table 1. ^26^ ^27^ ^5^ ^28^ ^29^ ^30^

From Figure 1 and Figure 2 one can see that the main differences between the mildly malignant and malignant brain tumors occur at 1585 cm^−1^ corresponding to reduced cytochrome c (Fe^+2^). ^28^

In order to understand better the difference between mildly malignant and malignant brain tumors we prepared the average Raman spectra, presented on Figure 3. The average spectra results are based on thousands of single spectra obtained from the Raman images. To determine the significant difference between the mildly malignant and malignant tumor tissue we present the differential spectrum (mildly malignant brain tissue subtracted from malignant brain tissue) in the fingerprint region and in the high-frequency region (Figure 3B).

**Figure 3.**
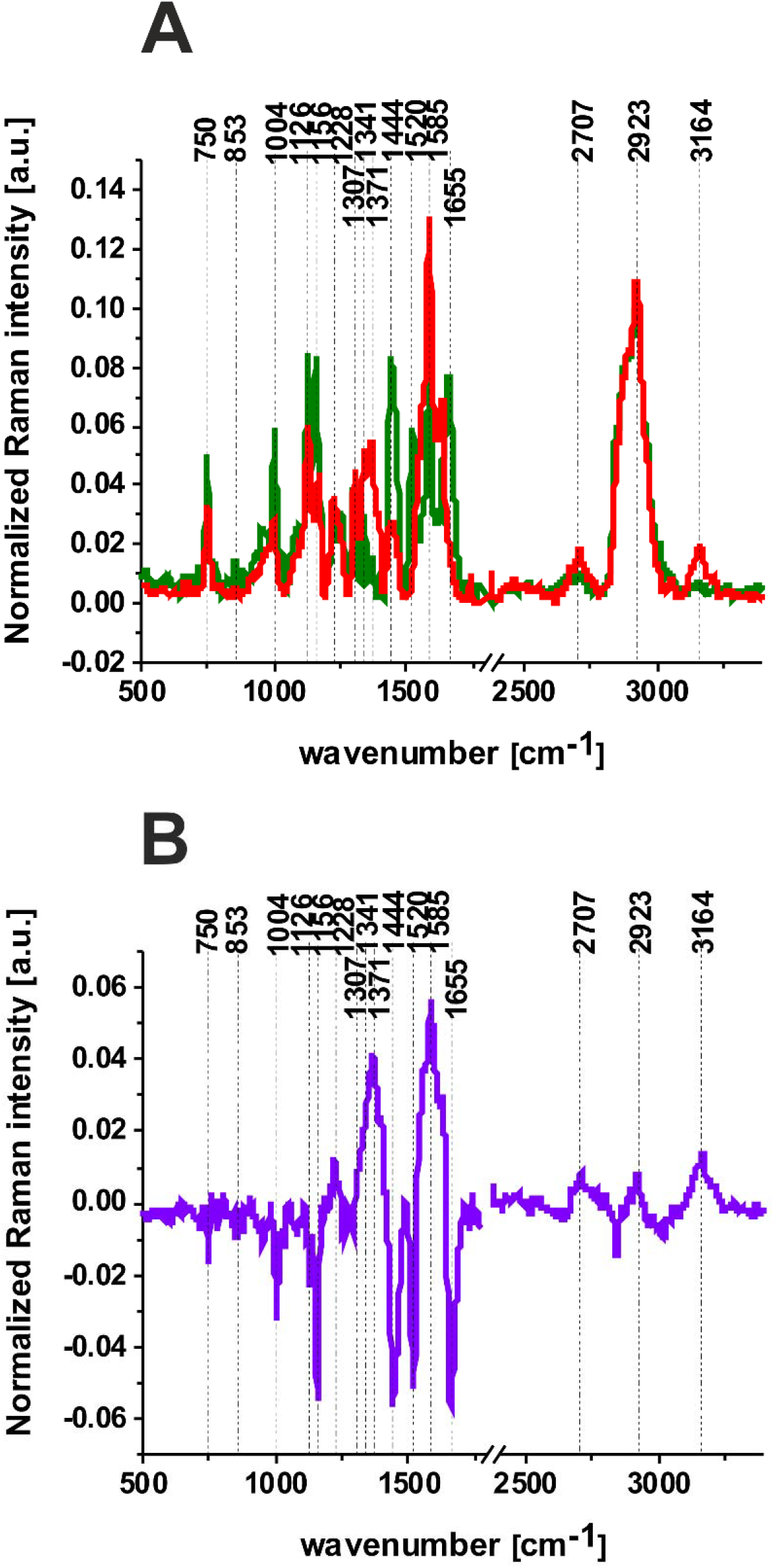
*Average Raman spectrum of meningothelioma WHO grade I (green line) and anaplastic oligodendroglioma WHO grade III (red line) in the fingerprint region and in the high-frequency region (A); differential spectrum (malignant tumor-mildly malignant tumor) (purple line) in the fingerprint region and in the high-frequency region (B).*

Comparing the spectra from Figure 3 more detailed biochemical information about the changes in cancer development is obtained. One can see that main biochemical differences are observed in the fingerprint region. Firstly, in malignant tumor the Raman intensities of the bands at 1307, 1341, 1371 and 1585 cm^−1^ are higher than in mildly malignant. Secondly the bands at 750, 1004, 1126, 1156, 1444 and 1655 cm^−1^ are much stronger in mildly malignant brain tissue than in malignant ones. We can see that the most prominent changes that occur due to the development of cancer are connected with the peak 1585 cm^−1^ assigned to the vibration of heme group of cytochrome c. ^28^ The Raman intensity (and concentration) of reduced cytochrome c increases with brain tumor aggressiveness. Previous studies reported similar observations. ^28^

Spectacular difference is also observed at 1444 cm^−1^. This band attributed to the lipids and fatty acids is much stronger in the mildly malignant tissue. Our results show higher lipid content in tumors of lower grade of malignancy. Previous studies reported similar observations. ^31^ ^32^ ^33^ ^34^

This study expands our previous research by adding lipid content as an additional signature of cancer, motivated by recent works that suggest, that abnormal lipid metabolism can be another hallmark for various cancers that can be useful in a stratification of malignancy. ^32^ ^31^ ^33^ ^34^

The correlation between lipid metabolism and cancer invasiveness is further explored in breast cancer. To test biochemical composition of human breast tissue we performed the same Raman procedure for invasive ductal carcinoma G3. Figure 4 presents the video image, Raman image and Raman spectra of the cancerous human breast tissue.

**Figure 4.**
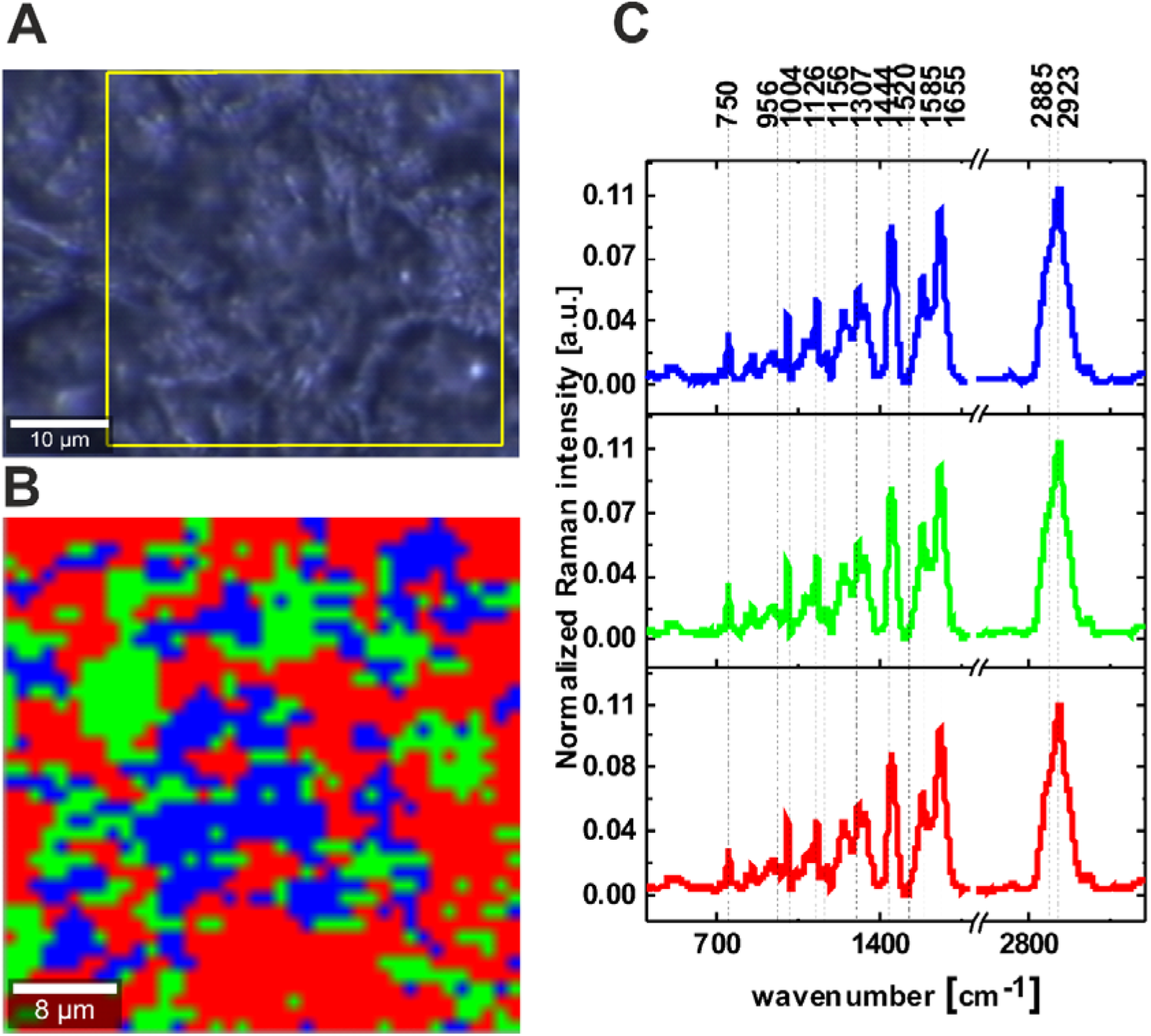
*Microscopy image (A), Raman image (40 µm x 40 µm) (B) and Raman spectra (C) obtained by Cluster Analysis methods in range 400-3300 cm^−1^ from invasive ductal carcinoma G3. Integration time for Raman images 0.5 s in the high frequency region and 0.5 s in the fingerprint region. The line colours of the Raman spectra correspond to the colours of Raman image*

One can see from Fig. 4 that cancerous breast tissue is dominated by peaks 750, 956, 1004, 1126, 1307, 1444, 1585, 1655, 2885 and 2923 cm^−1^. We do not observe Raman peaks at 1156 and 1520 cm^−1^ attributed to carotenoids that are clearly visible in mildly malignant tumor (Fig. 3).

To understand the biochemical changes that occur in tumors we measured and analyzed the Raman spectra in various types of human brain tumors and compared with breast cancer. Figure 5 shows the Raman spectra of the brain tumor tissue (testicular cancer metastasis, gliosarcoma WHO IV, anaplastic oligodendroglioma WHO III, meningioma WHO II, meningothelioma WHO I, pituitary adenoma, neurofibroma and invasive ductal carcinoma G3) in the fingerprint (A) and high frequency region (B).

**Figure 5.**
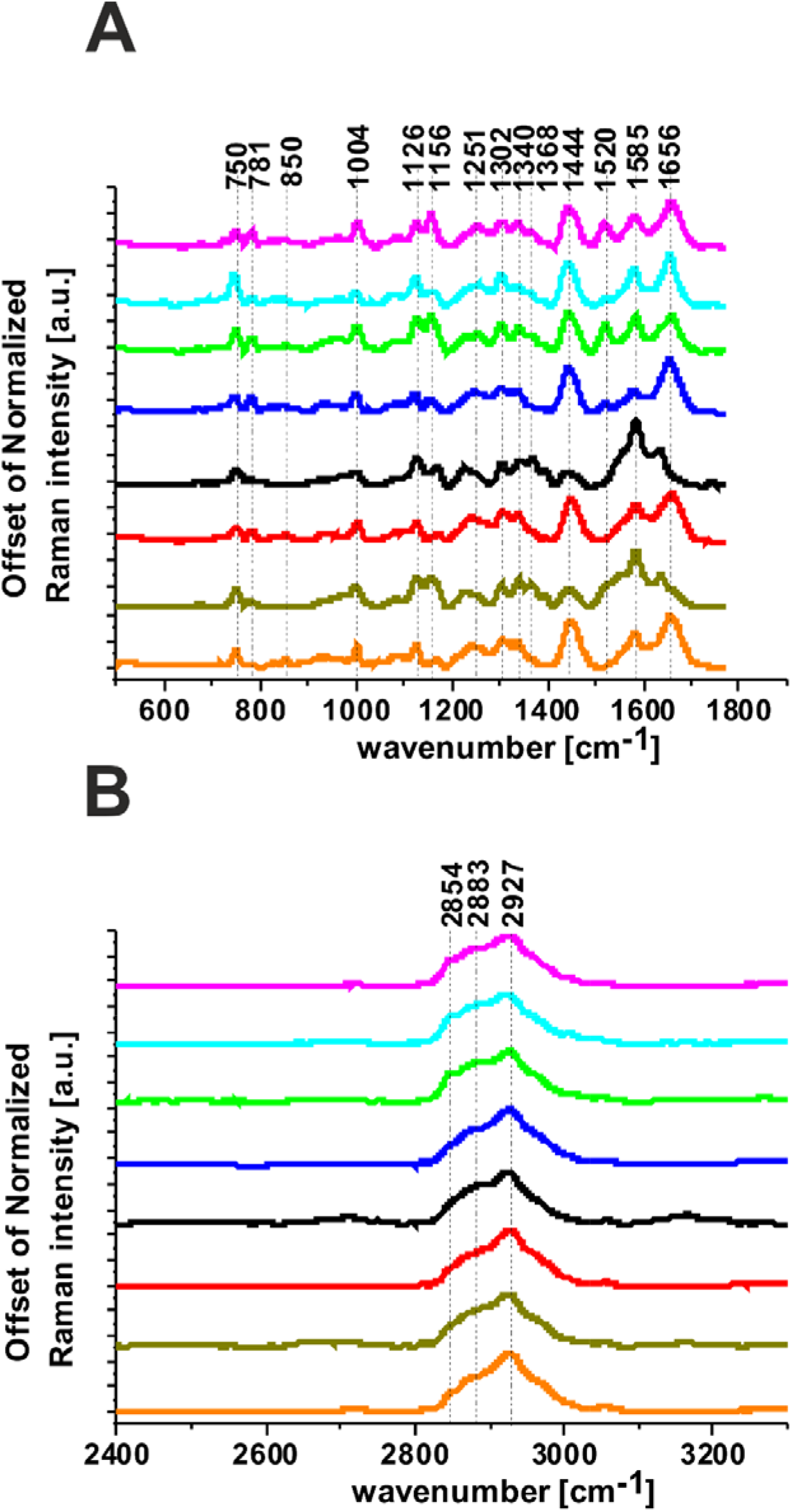
*The Raman spectra of the invasive ductal carcinoma G3 (orange), testicular cancer metastasis (olive), gliosarcoma WHO IV (red), anaplastic oligodendroglioma WHO III (black), meningioma WHO II (blue), meningothelioma WHO I (green), pituitary adenoma (turquoise) and neurofibroma (magenta) in the fingerprint (A) and high frequency region (B).*

Many significant biochemical differences between the various types of brain tumor can be observed. The most relevant differences presented on Fig. 5 are related to the vibrations of carotenoids (1156 cm^−1^, 1520 cm^−1^), fatty acids (1444 cm^−1^, 1655 cm^−1^), proteins (1004 cm^−1^, 1655 cm^−1^) and cytochrome (1585 cm^−1^). First we observe that the band 1444 cm^−1^ assigned to fatty acids is much stronger in all mildly malignant cancers. In contrast the band at 1585 cm^−1^ is much stronger in more aggressive brain tumor tissues. This band has a lower intensity in the mildly malignant brain tissue, indicating that the reduced cytochrome c concentration is much lower in the mildly malignant human brain tumor tissue than in the malignant tissue. Moreover, the bands of carotenoids (1156 cm^−1^ and 1520 cm^−1^) are more visible in mildly malignant brain tumors. The band at 1656 cm^−1^ corresponding to fatty acids and proteins has higher intensity in the mildly malignant brain tumor tissue. From high frequency region (Fig. 5B) one can see that the band at 2854 cm^−1^ assigned to lipids show higher lipid content in mildly malignant human brain tumor tissue.

To access the diagnostic potential of Raman spectroscopy in estimating the type of human brain tumor we calculated the Raman intensity ratios for characteristic Raman vibrations and obtained the ratios of the characteristic Raman peaks that can be useful in a stratification of malignancy for various cancers were selected.

Table 2 and Figure 6 illustrate the intensity and standard deviation for the ratios 1585/1655, 1585/1444, 1520/1585, 1156/1585, 1004/1585. Presented results are based on thousands of Raman spectra for each patient obtained from cluster analysis.

**Figure 6.**
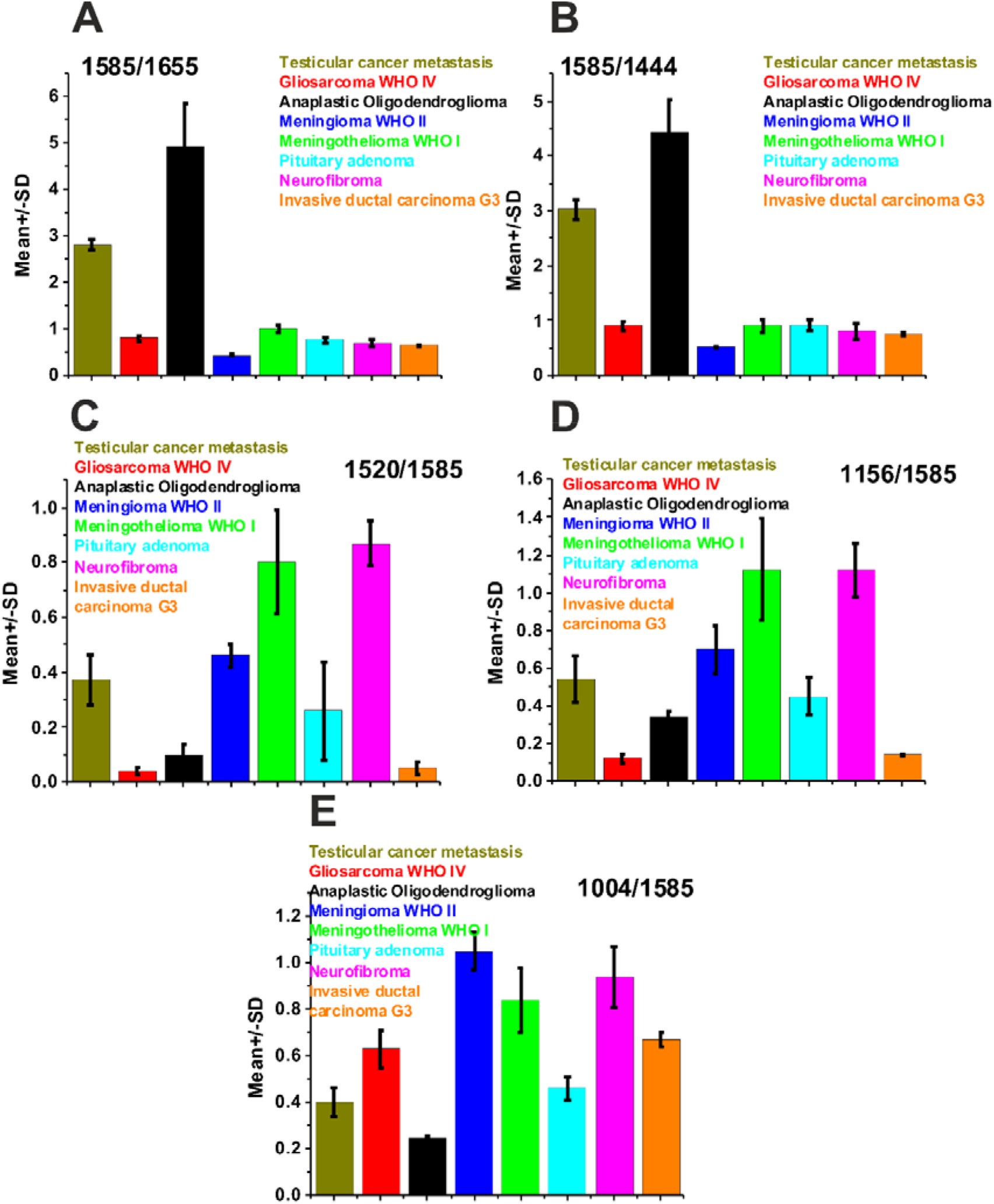
*The intensity ratios and SD for 1585/1655 (A), 1585/1444 (B), 1520/1585 (C), 1156/1585(D), 1004/1585 (E) for testicular cancer metastasis (olive), gliosarcoma WHO IV (red), anaplastic oligodendroglioma WHO III (black), meningioma WHO II (blue), meningothelioma WHO I (green), pituitary adenoma (turquoise), neurofibroma (magenta), invasive ductal carcinoma G3 (orange)*

**Table 2.** *Raman intensity ratios, 1585/1655, 1585/1444, 1520/1585, 1156/1585, 1004/1585 for, testicular cancer metastasis, gliosarcoma WHO IV, anaplastic oligodendroglioma WHO III, meningioma WHO II, meningothelioma WHO I, pituitary adenoma, neurofibroma, invasive ductal carcinoma G3, SD – standard deviation.*

| ratio | Testicular cancer<br>metastasis | gliosarcoma WHO<br>IV | Anaplastic<br>Oligodendroglioma<br>WHO III | meningioma WHO<br>II | meningothelioma<br>WHO I | Pituitary adenoma | Neurofibroma | Invasive ductal<br>carcinoma G3 |
| --- | --- | --- | --- | --- | --- | --- | --- | --- |
|  | Mean± SD | Mean± SD | Mean±SD | Mean± SD | Mean±SD | Mean± SD | Mean±SD | Mean±SD |
| 1585/<br>1655 | 2.8±0.11 | 0.8±0.06 | 4.95±0.89 | 0.44±0.02 | 1.02±0.09 | 0.76±0.06 | 0.7±0.08 | 0.65±0.02 |
| 1585/<br>1444 | 3.04±0.18 | 0.9±0.07 | 4.43±0.57 | 0.52±0.02 | 0.9±0.11 | 0.92±0.09 | 0.81±0.14 | 0.74±0.03 |
| 1520/<br>1585 | 0.37±0.09 | 0.04±0.01 | 0.1±0.04 | 0.46±0.04 | 0.8±0.19 | 0.26±0.18 | 0.87±0.08 | 0.05±0.02 |
| 1156/<br>1585 | 0.54±0.12 | 0.12±0.02 | 0.34±0.03 | 0.7±0.13 | 1.12±0.27 | 0.45±0.1 | 1.12±0.14 | 0.14±0.006 |
| 1004/<br>1585 | 0.4±0.06 | 0.63±0.08 | 0.25±0.01 | 1.05±0.08 | 0.84±0.14 | 0.46±0.05 | 0.94±0.13 | 0.67±0.03 |

One can see from Figure 6 and Table 2 that presented above ratios are useful to discriminate the type of human brain tumor. Analyzing the ratios for vibrations of lipids and proteins non-negligible differences are visible for the less aggressive brain tumor in comparison to the malignant brain tumor. The ratios 1585/1655 and 1585/1444 are significantly higher for malignant brain tumor tissue (testicular cancer metastasis, anaplastic oligodendroglioma WHO III). While for less aggressive human brain tissue (meningothelioma WHO I, meningioma WHO II, pituitary adenoma and neurofibroma) the ratios of 1520/1585, 1156/1585 and 1004/1585 are higher. All values of ratio presented on Fig. 6 for invasive breast carcinoma are comparable to the ratios for the gliosarcoma WHO IV. The observed discrimination between the various types of human brain tumor and breast cancer is based on Raman intensity ratios.

To better visualize the differences between various human brain tumors and human breast cancer we used chemometric analysis by statistical analysis using PCA and PLS-DA methods.

Figure 7 shows the PCA and PLS-DA score plot for the Raman spectra of human brain and breast cancers. Figure 7 shows evidently that differences between the cancer subtypes can be revealed.

**Figure 7.**
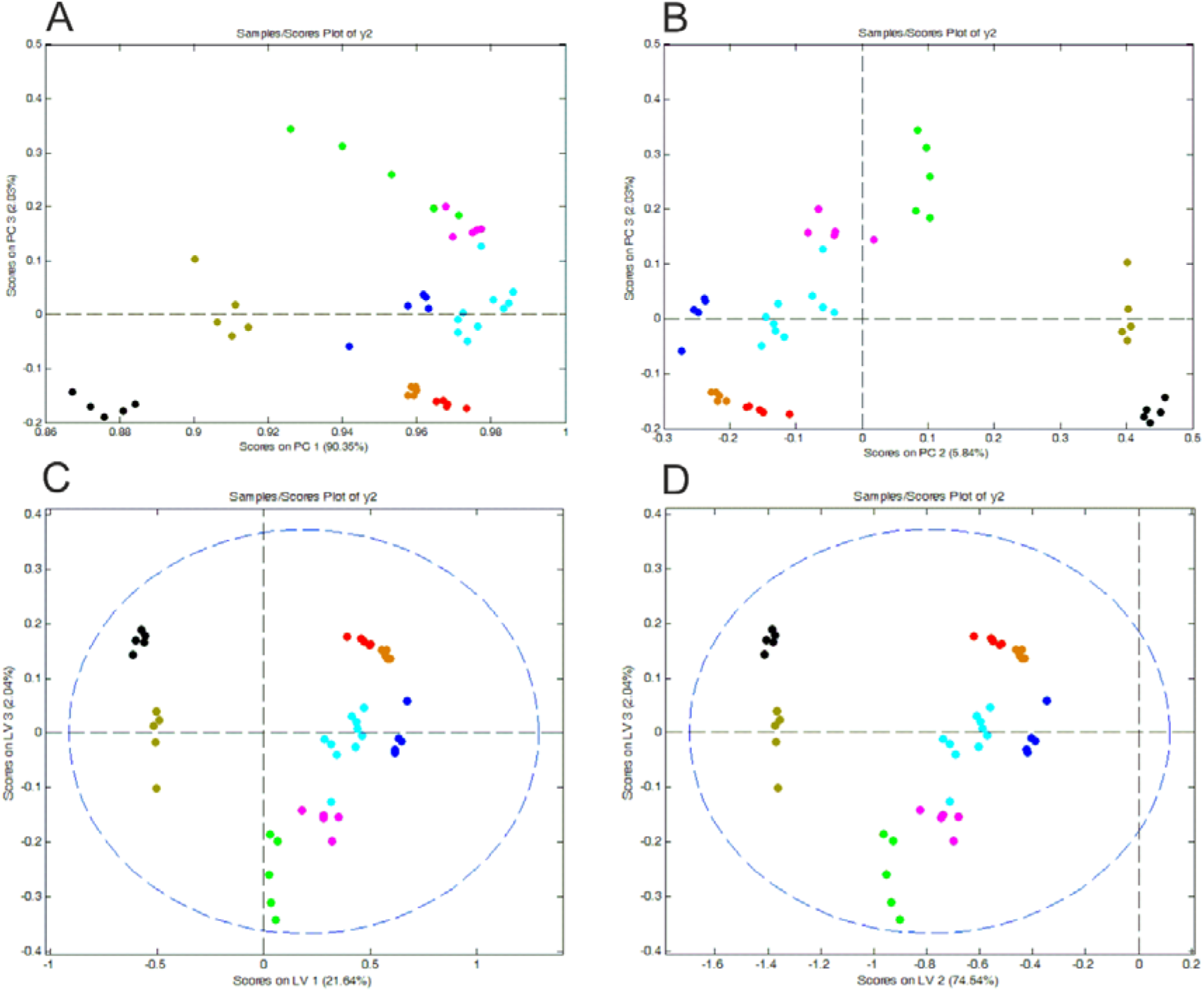
*PCA (A, B) and PLSDA (C, D) score plot for the average Raman spectra for: Testicular cancer metastasis (olive), gliosarcoma WHO IV (red), anaplastic oligodendroglioma WHO III (black), meningioma WHO II (blue), meningothelioma WHO I (green), pituitary adenoma (turquoise), neurofibroma (magenta), invasive ductal carcinoma G3 (orange)*

The plots presented on Figure 7 confirm the differences between human brain and breast cancers. The differences and similarities are visible by grouping the results into separate clusters. Moreover, Raman spectra for the most aggressive brain tumor (anaplastic oligodendroglioma WHO III (black colour) and testicular cancer metastasis (olive colour)) are the most separated from the rest of cancer subtypes. One can see that the Raman spectra for the most aggressive brain tumor (gliosarcoma WHO IV (red colour)) are mixed with Raman spectra for the aggressive breast cancer G3 (orange colour). This result was presented also on Fig. 6 and Table 2.

To access the diagnostic potential of Raman spectroscopy and imaging for clinical practice we calculated sensitivity and specificity. Table 3 presents the value of sensitivity and specificity obtained from PLS-DA method. The high values for sensitivity and specificity highlight the importance of Raman spectroscopy and imaging as a new diagnostic tool.

**Table 3.** *The value of sensitivity and specificity for calibration and cross validation procedure from PLS-DA analysis.*

|  | Testicular cancer<br>metastasis | gliosarcoma WHO<br>IV | Anaplastic<br>Oligodendroglioma<br>WHO III | meningioma WHO II | meningothelioma<br>WHO I | Pituitary adenoma | Neurofibroma | Invasive ductal<br>carcinoma G3 |
| --- | --- | --- | --- | --- | --- | --- | --- | --- |
| Sensitivity (calibration) | 1.0 | 1.0 | 1.0 | 1.0 | 1.0 | 1.0 | 1.0 | 1.0 |
| Specificity (calibration) | 0.875 | 0.875 | 1.0 | 1.0 | 0.815 | 0.457 | 0.850 | 0.9 |
| Sensitivity (cross validation) | 1.0 | 0.8 | 1.0 | 1.0 | 1.0 | 0.9 | 1.0 | 1.0 |
| Specificity (cross validation) | 0.875 | 0.9 | 1.0 | 1.0 | 0.9 | 0.486 | 0.850 | 0.9 |

## Conclusions

In this exploratory study, we demonstrated that Raman spectroscopy, Raman imaging and statistical analysis are tools for human cancer diagnostics. For human brain tumor distinguishing vibrational signatures were primarily responsible for alterations in carotenoids, proteins, fatty acids and cytochrome c. Presented research gave the opportunity to determine Raman biomarkers differentiating various types of human brain tumor. We have shown that bands characteristic for carotenoids, proteins, fatty acids and cytochrome c can be used as universal biomarkers to distinguish aggressiveness in human brain tumors. The ratios 1585/1655, 1585/1444, 1520/1585, 1156/1585, 1004/1585 can be used as a Raman biomarkers for diagnosing human brain tumors. We have demonstrated that Raman spectroscopy combined with statistical analysis would be nonsubjective technique used as a method alongside neuropathology for brain cancer diagnosis.

## Materials and Methods

### Tissue preparation

The brain tissues were obtained from the material removed over the operation from Department of Neurosurgery, Spine and Peripheral Nerve Surgery, University Hospital WAM-CSW, Lodz. The breast tissue was obtained during routine surgery from the WWCOiT Nicolaus Copernicus in Lodz. Obtaining samples did not affects the course of the operation or treatment of the patients. The tissue brain and breast sections after cutting into 16 µm thick slices in microtome were put on CaF_2_ windows (Crystran). For all experiments we used only human fresh brain and breast tissue. We do not use the procedure formalin fixation, paraffin-embedding because of its influence on Raman measurements. ^35^ Written informed consent was obtained from patients. All experiments were conducted in accordance with relevant guidelines and regulations of the Bioethical Committee at the Medical University of Lodz, Poland (RNN/247/19/KE) and (RNN/323/17/KE/17/10/2017).

### Raman spectroscopy, Raman imaging and statistical methods

All Raman spectra and Raman imaging presented in this manuscript were obtained using alpha 300 RSA+ (WITec, Germany) combined with confocal microscope coupled via the fibre of a 50 mm core diameter with an UHTS (Ultra High Throughput Spectrometer) spectrometer and a CCD Camera (Andor Newton DU970N-UVB-353) operating in standard mode with 1600×200 pixels at 60C with full vertical binning. All experiment were performed using laser beam (SHG of the Nd:YAG laser (532 nm)) and 40x dry objective (Nikon, objective type CFI Plan Fluor CELWD DIC-M, numerical aperture (NA) of 0.60 and a 3.6–2.8 mm working distance). All Raman experiments were performed using laser with a power 10 mW at the sample position. Every day before Raman measurements the confocal system was calibrated using silicon plate (520.7◻cm^−1^). Each Raman spectra and Raman imaging were processed to remove cosmic rays, smoothing (Savitzky-Golay method) and background subtraction. Preprocessing was performed using WITec Plus Software. Raman maps data were analysed using Cluster Analysis method. Detailed description of equipment and methodology used in the paper is available elsewhere ^31^ ^36^ ^37^

The Raman spectra were obtained from 9 patients. Among patients with brain tumors 1 were diagnosed with testicular cancer metastasis, 1 with gliosarcoma WHO IV, 1 with anaplastic oligodendroglioma WHO III, 1 with meningioma WHO II, 1 with meningothelioma WHO I, 2 with pituitary adenoma and 1 with neurofibroma. Breast cancer sample was diagnosed with invasive ductal carcinoma G3. For each patient thousands of single spectra from different sites of the sample were obtained from cluster analysis. In detail, for sample with testicular cancer metastasis we used typically 8400 Raman spectra for averaging, for sample with gliosarcoma WHO IV 6400 Raman spectra, for sample with anaplastic oligodendroglioma WHO III 9200 Raman spectra, for sample with meningioma WHO II 3000 Raman spectra, for sample with meningothelioma WHO I 90000 Raman spectra, for 2 patients with pituitary adenoma 15000 Raman spectra, for sample with neurofibroma 3600 Raman spectra and for the sample with invasive ductal carcinoma G3 1600 Raman spectra, respectively. Principal Component Analysis and Partial Least Squares Discriminant Analysis were performed using Mathlab and PLS_Toolbox Version 4.0. Detailed description of chemometric methods was presented in our previous paper ^38^ ^31^. A significance level of less than 0.05 was used for all statistical analyses.

## Acknowledgement

This work was supported by the National Science Centre of Poland (Narodowe Centrum Nauki, UMO-2019/33/B/ST4/01961 and Miniatura 4 (grant 2020/04/X/ST4/00325)).

## Notes

### Competing Interest Statement

The authors have declared no competing interest.

## References

1 C. Hardwidge and S. Hettige, Surgery, 2012, 30, 155–161.

2 F. K. Lu, D. Calligaris, O. I. Olubiyi, I. Norton, W. Yang, S. Santagata, X. S. Xie, A. J. Golby and N. Y. R. Agar, Cancer Res., 2016, 76, 3451–3462.

3 M. Jermyn, J. Desroches, J. Mercier, K. St-Arnaud, M.-C. Guiot, F. Leblond and K. Petrecca, Biomed. Opt. Express, 2016, 7, 5129.

4 D. Bovenkamp, A. Micko, J. Püls, F. Placzek, R. Höftberger, G. Vila, R. Leitgeb, W. Drexler, M. Andreana, S. Wolfsberge and A. Unterhuber, Molecules, 2019, 24, 1–15.

5 L. M. Fullwood, G. Clemens, D. Griffiths, K. Ashton, T. P. Dawson, R. W. Lea, C. Davis, F. Bonnier, H. J. Byrne and M. J. Baker, Anal. Methods, 2014, 6, 3948–3961.

6 S. Portet, R. Naoufal, G. Tachon, A. Simonneau, A. Chalant, A. Naar, S. Milin, B. Bataille and L. Karayan-Tapon, Neuro-Oncology Adv., 2019, 1, 1–12.

7 D. N. Louis, A. Perry, G. Reifenberger, A. von Deimling, D. Figarella-Branger, W. K. Cavenee, H. Ohgaki, O. D. Wiestler, P. Kleihues and D. W. Ellison, Acta Neuropathol., 2016, 131, 803–820.

8 C. L. M. Morais, T. Lilo, K. M. Ashton, C. Davis, T. P. Dawson, N. Gurusinghe and F. L. Martin, Analyst, 2019, 144, 7024–7031.

9 V. K. Puduvalli, M. Hashmi, L. D. McAllister, V. A. Levin, K. R. Hess, M. Prados, K. A. Jaeckle, W. K. A. Yung, S. S. Buys, J. M. Bruner, J. J. Townsend, R. Davis, R. Sawaya and A. P. Kyritsis, Oncology, 2003, 65, 259–266.

10 R. Stupp, W. P. Mason, M. J. Van Den Bent, M. Weller, B. Fisher, M. J. B. Taphoorn, K. Belanger, A. A. Brandes, C. Marosi, U. Bogdahn, J. Curschmann, R. C. Janzer, S. K. Ludwin, T. Gorlia, A. Allgeier, D. Lacombe, J. G. Cairncross, E. Eisenhauer and R. O. Mirimanoff, N Engl J Med, 2005, 352, 987–96.

11 P. Mulvenna, M. Nankivell, R. Barton, C. Faivre-Finn, P. Wilson, E. McColl, B. Moore, I. Brisbane, D. Ardron, T. Holt, S. Morgan, C. Lee, K. Waite, N. Bayman, C. Pugh, B. Sydes, R. Stephens, M. K. Parmar and R. E. Langley, Lancet, 2016, 388, 2004–2014.

12 T. R. Daniels, E. Bernabeu, J. A. Rodríguez, S. Patel, M. Kozman, D. A. Chiappetta, E. Holler, J. Y. Ljubimova, G. Helguera and M. L. Penichet, Biochim. Biophys. Acta - Gen. Subj., 2012, 1820, 291–317.

13 M. C. Chamberlain and S. Johnston, Cancer, 2009, 115, 1734–1743.

14 Y. Zhou, C.-H. Liu, B. Wu, X. Yu, G. Cheng, K. Zhu, K. Wang, C. Zhang, M. Zhao, R. Zong, L. Zhang, L. Shi and R. R. Alfano, J. Biomed. Opt., 2019, 24, 1.

15 J.R. Antonio, E.M. Goloni-Bertollo and L.A. Tridico, An Bras Dermatol, 2013, 88, 329–343.

16 D.J. Canale and J. Bebin Von Recklinghausen disease of the nervous system. In: Vinken4.PJ, Bruyn GW, editors. Handbook of clinical neurology. Amsterdam: Holland Publishing Company; 1972, 14, 132–162.

17 K. Khosrotehrani, S. Bastuji-Garin, J. Zeller, J. Revuz and P. Wolkenstein, Arch Dermatol. 2003, 139, 187–191.

18 M.E. Molitch, C. Review & Education, 2017, 317, 6–11.

19 A. Willett, J. Ben Wilkinson, C. Shah and M. P. Mehta, Indian J. Med. Paediatr. Oncol., 2015, 36, 87–93.

20 K. Tashibu, No To Shinkei, 1990, 42, 999–1004.

21 D. T. Theodosis, D. A. Poulain and S. H. R. Oliet, Physiol. Rev., 2008, 88, 983–1008.

22 H. Abramczyk, A. Imiela, B. Brożek-Płuska, M. Kopeć, J. Surmacki and A. Śliwińska, Cancers, 2019, 11, 1–25.

23 J. Mill, T. Tang, Z. Kaminsky, T. Khare, S. Yazdanpanah, L. Bouchard, P. Jia, A. Assadzadeh, J. Flanagan, A. Schumacher, S. C. Wang and A. Petronis, Am. J. Hum. Genet., 2008, 82, 696–711.

24 J. J. Miller and K. G. Baimbridge, Brain Res., 1983, 278, 322–326.

25 O. Uckermann, W. Yao, T. A. Juratli, R. Galli, E. Leipnitz, M. Meinhardt, E. Koch, G. Schackert, G. Steiner and M. Kirsch, J. Neurooncol., 2018, 139, 261–268.

26 H. Abramczyk and A. Imiela, Spectrochim. Acta - Part A Mol. Biomol. Spectrosc., 2018, 188, 8–19.

27 K. Gajjar, L. D. Heppenstall, W. Pang, K. M. Ashton, J. Trevisan, I. I. Patel, V. Llabjani, H. F. Stringfellow, P. L. Martin-Hirsch, T. Dawson and F. L. Martin, Anal. Methods, 2013, 5, 89–102.

28 H. Abramczyk, B. Brozek-Pluska, M. Kopec, J. Surmacki, M. Blaszczyk, M. Radek, Cancers, 2021, 13, 1–20.

29 H. Abramczyk, J. M. Surmacki and B. Brozek-Pluska, bioRxiv, DOI:10.1101/2020.05.08.083915.

30 M. Gasior-Głogowska, M. Komorowska, J. Hanuza, M. Maczka and M. Kobielarz, Acta Bioeng. Biomech., 2010, 12, 53–60.

31 J. Surmacki, B. Brozek-Pluska, R. Kordek and H. Abramczyk, Analyst, 2015, 140, 2121–2133.

32 C. Nieva, M. Marro, N. Santana-Codina, S. Rao, D. Petrov and A. Sierra, PLoS One, 2012, 10, 1–10.

33 H. Abramczyk, B. Brozek-Pluska, J. Surmacki, J. Jablonska-Gajewicz and R. Kordek, Prog. Biophys. Mol. Biol., 2012, 108, 74–81.

34 H. Abramczyk, J. Surmacki, M. Kopeć, A. K. Olejnik, K. Lubecka-Pietruszewska and K. Fabianowska-Majewska, Analyst, 2015, 140, 2224–2235.

35 B. Brozek-Pluska, M. Kopec, J. Surmacki and H. Abramczyk, Infrared Phys. Technol., 2018, 93, 247–254.

36 M. Kopeć and H. Abramczyk, Spectrochim. Acta - Part A Mol. Biomol. Spectrosc., 2018, 198, 338–345.

37 H. Abramczyk and B. Brozek-Pluska, Anal. Chim. Acta, 2016, 909, 91–100.

38 B. Brozek-Pluska, M. Kopeć and H. Abramczyk, Anal. Methods, 2016, 8, 8542–8553.

